# Movement-integrated habitat selection reveals wolves balance ease of travel with human avoidance in a risk-reward trade-off

**DOI:** 10.1101/2024.01.04.574266

**Authors:** Katrien A. Kingdon, Christina M. Prokopenko, Daniel L. J. Dupont, Julie Turner, Jonathan Wiens, Vanessa B. Harriman, Eric Vander Wal

**Affiliations:** Memorial University of Newfoundland, Department of Biology, 45 Arctic Avenue, St. John’s, NL Canada A1C 5S7; Memorial University of Newfoundland, Cognitive and Behavioural Ecology Interdisciplinary Program, 232 Elizabeth Avenue, St. John’s, NL Canada A1B 3X9; Université de Saint-Boniface, Sciences expérimentales, 200, avenue de la Cathédrale, Winnipeg, MB Canada R2H 0H7; Manitoba Hydro, Transmission & Distribution Environment and Engagement Department, 360 Portage Avenue, Winnipeg, MB Canada R3C 0G8; Ducks Unlimited Canada, Institute for Wetland and Waterfowl Research, 1 Mallard Bay, Stonewall, MB Canada R0C 2Z0

**Keywords:** animal movement, *Canis lupus*, resource selection, roads, step selection, space use, transmission lines

## Abstract

Anthropogenic linear features are often linked to alterations in wildlife behaviour and movement. Landscape features such as habitat can have important mediating effects on wildlife response to disturbance and yet is rarely explicitly considered in how it interacts with other features. This study tests the effects of habitat variation on the space-use responses of GPS-collared wolves to linear features in eastern Manitoba. We simultaneously model wolf movement and selection within a conditional logistic regression framework (integrated Step Selection Analysis) with explicit consideration for how habitat alters these responses through either added cover or friction to movement. Classifying linear features based on the selection and movement response of wolves revealed that pairing transmission lines and primary roads increased the avoidance response to be greater than either feature on their own and provided evidence of a semi-permeable barrier to movement. In contrast, features with reduced human activity, including secondary and tertiary roads were highly selected for and provided access to movement corridors. Explicit parameterization of habitat provides evidence that where a linear feature is routed and which habitats it interacts with will have the greatest implications for wolf behavioural responses. Reduction of avoidance behavioural in highly risky environments signifies the importance of habitat for maintaining landscape connectivity, particularly when routing multiple features together. Natural regrowth along these features reduced movement advantages by increasing friction, indicating that actively decommissioning other features such as secondary roads would have important implications for reducing wolf encounters with prey. Knowing the influence of adjacent habitats on the likelihood of wolves selecting for a given linear feature creates context to minimize the impact of new anthropogenic features on behaviour.

## Introduction

Rapid anthropogenic alteration of natural landscapes is challenging the stability of many ecosystems globally (DeCesare, 2012; Ryall and Fahrig, 2006). Anthropogenic linear features produce long, cleared sections across a landscape (Knight and Kawashima, 1993) and are the cause of much habitat loss and fragmentation (Luque et al., 2013). For wildlife, landscape changes create a tension between resource acquisition and avoidance of areas associated with humans (Ripple and Beschta, 2012). This trade-off between risk and reward for animals shapes animal behaviour including movement patterns and resource selection (Coulson et al., 2011; Frid and Dill, 2002; Sih et al., 2011). Linear features can be considered high-risk if the feature is associated with areas of high human activity, such as primary roads, where wildlife are more likely to interact with humans. Features with reduced human use, however, such as hydroelectric right-of-ways (ROWs), are low-risk for wildlife. The surrounding habitat can become a mediating factor in how wildlife interact with linear features, where individuals alter their habitat selection to gain food or safety (Hugie and Dill, 1994; Sih et al., 1998) or increase their ease of movement. Changes in wildlife behaviour and movement can have cascading effects through the ecosystem, influencing both intraspecific and interspecific interactions (Schmitz et al., 1997). Understanding how wildlife populations experience different trade-offs along high and low-risk linear features can provide critical insight into animal ecology, conservation, and management.

Consequences of linear features on the landscape for wildlife can be direct, such as increased collisions with vehicles along roads, and indirect when wildlife are displaced to avoid human activity (Neumann et al., 2012; Polfus et al., 2011). The high level of human activity associated with some linear features often discourages wildlife, establishing barriers to wildlife movement (Beyer et al., 2016; Latham et al., 2011). Woodland caribou (*Rangifer tarandus caribou*) cross roads up to six times less frequently than expected from random simulations (Dyer et al., 2002) and elk (*Cervus elaphu*s) avoid roads across multiple scales of winter range selection (Christina M Prokopenko et al., 2017). Further, some grizzly bear populations experience a genetic discontinuity at highways, indicating that the highways impose a physical barrier to movement with population level consequences (Sawaya et al., 2014). Other features, such as hydroelectric ROWs, experience little to no human activity but still influence movements and distributions (Courbin et al., 2014; Semeniuk et al., 2014). For example, both ravens (*Corvus corax*) and red-tailed hawks (*Buteo jamaicensis*) are more likely to nest along transmission lines than along roads or control transects (Knight and Kawashima, 1993). North American deermice (*Peromyscus maniculatus*) will readily cross transmission lines, while southern red-backed voles (*Myodes gapperi*) will instead move in parallel with the lines but avoid crossing them (Storm and Choate, 2012). Ultimately, the influence of linear features on a species will largely depend on how individuals use the landscape for resource acquisition and survival.

Predator species are particularly susceptible to anthropogenic environmental change, as they are wide-ranging and have high metabolic requirements, two traits that result in high rates of human-wildlife conflict, and conflict-related wildlife mortality (Ripple and Beschta, 2012). For example, canids that increase avoidance of roads during the day are more likely to survive than individuals that do not adjust their selection (Benson et al., 2015). In addition, linear features can influence the use of the surrounding landscape where, for example, roads have reduced the quality of the surrounding habitat for wolverines (*Gulo gulo)* (Scrafford et al., 2018). Alternatively, linear features such as hydroelectric ROWs and logging roads may promote predator use. Increased use of a linear feature can consequently enhance predation rates on prey by increasing predator abundance along the linear feature (Andrén, 1992), facilitating predator hunting efficiency (McKenzie et al., 2012), or increasing access to prey habitat through expanding patch edges (Ryall and Fahrig, 2006). Highly mobile species, including predators, likely encounter a wide range of linear features within their ranges; thus, the response to these environmental changes often leads to changes to encounter and kill rates with prey species (Fortin et al., 2005; Vander Vennen et al., 2016).

Gray wolves (*Canis lupus*) are apex predators of the boreal ecosystems, with their wide range converging with linear feature creation. Linear features can indirectly increase hunting efficiency through increased wolf movement linked to higher encounter rates with prey such as moose (Moffatt, 2012; Vander Vennen et al., 2016). Wolves will also combine selection for linear features and prey-preferred habitat to aid in their search (Lesmerises et al., 2012). Although wolves and other carnivores typically avoid areas with high anthropogenic activities (Gurarie et al., 2011; Whittington et al., 2005), expanding human populations and increasing demands for resource extraction will make linear features an increasingly present landscape feature. For gray wolves, a high density of primary roads within a territory can lead to constrained home range use (Gurarie et al., 2011). Additionally, wolves move at increased rates and in more linear directions through areas with high densities of linear features (Ehlers et al., 2014; Lesmerises et al., 2012). In contrast, wolves are more likely to select areas near trails with low levels of human use and are more likely to stay on them for longer periods of time (Whittington et al., 2005). Wolves also have been shown to travel faster on linear features such as seismic lines (McKenzie et al., 2012). Although human activity is a large driver of the risk-avoidance for wildlife, linear features can confer a movement advantage, particularly when the feature is surrounded by high-friction habitat such as dense forest or deep snow. Selection for linear features may therefore be driven by a balance between human risk and movement reward.

We test the risk-reward trade-offs between linear feature types and how these trade-offs influence gray wolf space use. Specifically, we examine how (1) wolf selection and crossing of linear features differ between those perceived to be high-risk compared to low-risk, as well as natural features such as waterways. (2) We measure how selection or avoidance of linear features influence movement rates either through increased ease of movement or increased vigilance when near them. (3) We compare linear feature selection between three common habitat types, coniferous, deciduous, and mixed, that differ in friction to movement. We hypothesize that wolf selection and avoidance will vary in response to the assumed level of human use (i.e., level of risk) associated with a given linear feature and that selection for linear features will increase when in habitats that increase friction to movement (i.e., level of reward). Consistent with the disturbance-risk hypothesis (Frid and Dill, 2002), we predict wolves will increase selection of areas near low-risk linear features and will cross them at higher rates in comparison to either high-risk features or natural features (P1). Additionally, we predict that when wolves are near or crossing linear features there will be an associated increase in movement rates, particularly when selecting for low-risk features (P2). Lastly, we predict that wolf selection and crossing of linear features will increase in dense habitats where ease of movement is decreased (P3).

## Methods

### Study Area

We would like to acknowledge this study takes place in Treaty 3 and Treaty 5 territories and in the homeland of the Métis Nation.

Our study was conducted in an ~7,200 km^2^ provincial management unit (Game Hunting Area 26, hereafter GHA 26), located in southeastern Manitoba, Canada. GHA 26 is bordered by Lake Winnipeg to the west and Ontario to the east (Figure 1A) and is part of the Boreal Shield Ecozone (Ecological Stratification Working Group 1995). The landscape in GHA 26 is composed predominantly of coniferous and mixed forests, interspersed with rock outcrops, rivers, lakes, and bogs. The area is predominantly public land, including Nopiming and Manigotagan River Provincial Parks, but also includes several communities. Resource extraction, including past forestry activity, has led to a network of roads, including primary roads, logging roads, and trails, throughout GHA 26. The additional presence of existing hydroelectric transmission ROWs that transect the area, and the construction of a new transmission ROW (2014-2016) are linear features that can impact wildlife movement.

**Figure 1.**
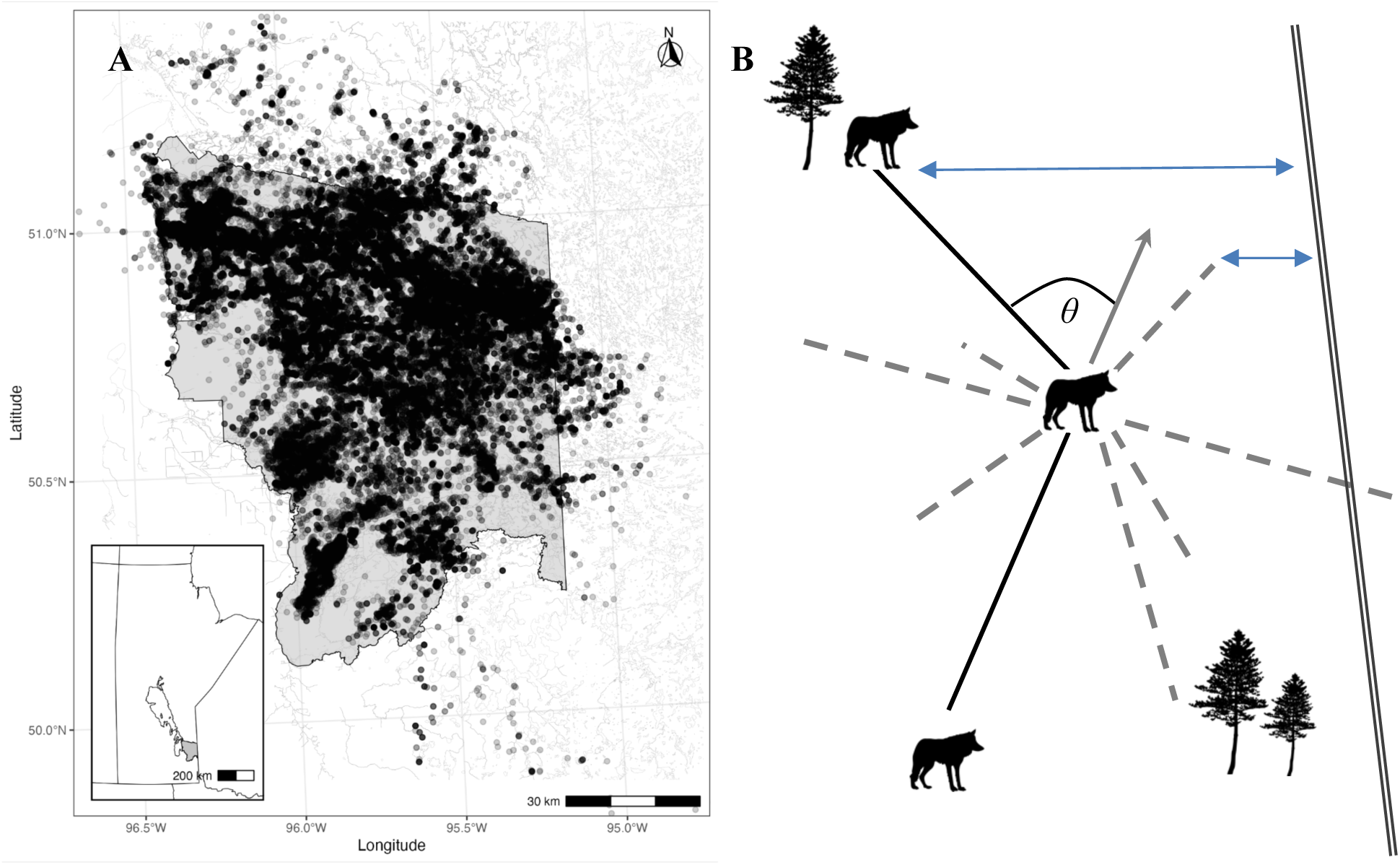
(A) Study area, Game Hunting Area 26, located in southeastern Manitoba, Canada, including relocations for all wolves (n=50) between 2014-2019; (B) Subset of relocations inferring movement (step length, turn angle) and selection (habitat, distance to a linear feature) through comparison of used steps (black lines) to available steps (grey).

### Collar Deployment

We captured and collared 46 wolves across 12 packs (minimum two collars per pack) between 2014-2019. The collared individuals overlapped with varying types of features including hydroelectric ROWs, roads, and natural linear features, such as waterways. Wolf captures were conducted by Bighorn Helicopters with a net-gun and occurred in accordance with the Memorial University Animal Care protocol 16-02-EV. Each wolf was fit with a GPS telemetry collar (Lotek Iridium TrackM 2D, Lotek Wireless Inc, Newmarket, ON, Canada; Sirtrack Pinnacle G5C, Sirtrack Limited, Hawkes Bay, New Zealand; Followit Tellus Medium 2D, Followit Sweden AB, Lindesberg, Sweden), which we programmed to collect a GPS relocation every two hours during all seasons.

### Integrated Step Selection Analysis

Common approaches to predicting animal space use patterns include habitat selection analyses (HSAs; e.g., resource selection analysis or RSA), which model the intensity an animal uses an available spatial unit as a function of both the availability (the area an animal could potentially use) and value of a given resource unit (Manly, 1974). However, movement and selection are intrinsically linked, both in the relation to the landscape and within the animal (Avgar et al., 2016). Movement-informed HSAs, including step selection analysis (SSA; Fortin et al. 2005) and integrated step selection analysis (iSSA; Avgar et al. 2016) are used to predict how environmental factors influence movement and selection (Figure 1B). In contrast to RSAs and SSAs, an iSSA defines availability based on probability distributions that are fit to observed movement behaviour including steps (linear connection between consecutive relocations) and turn angles (angular deviation in radians between consecutive relocations; Avgar et al. 2016).

iSSA determines selection by contrasting the steps used by an animal (hereafter, used steps) with steps that are available to the animal based on their observed movement behaviour (hereafter, available steps) (Avgar et al., 2016; Fortin et al., 2005). We randomly generated step lengths and turn angles from observed distributions of individual-level movement behaviour. We sampled available step lengths from a gamma distribution (population mean shape = 0.372, scale = 2844.4) of used steps demonstrating that short steps are taken more often with occasional longer steps. Available turn angles were sampled from values between π and −π corresponding to a Von Mises distribution where there is directional persistence based on either the previous or subsequent heading. For each used step, ten available steps were randomly generated from the population step length and turn angle distributions. Tracks, random steps, and covariates were extracted using the ‘amt’ package (Signer et al., 2019) in R v. 3.6.2 (R Core Team 2019). These available steps were then compared to used points to determine selection and movement behaviours. We fit our iSSA model using the ‘glmmTMB’ package (Brooks et al., 2017) using a mixed effects model approach with a Poisson error distribution (Muff et al., 2020).

### Core Model Covariates

A core model that included covariates expected to influence movement and selection regardless of proximity to a linear feature was used as a foundation in each subsequent model. We included four covariates in the core model including step length, turn angle, habitat, and time of day. Step lengths were transformed using the natural logarithm, which modifies the shape parameter of the gamma distribution. We extracted the cosine of all turn angles to transform the circular measure into a linear indicator where a value of −1 indicates a complete reversal in direction and 1 corresponds to linear persistence (Avgar et al., 2016; Prokopenko et al., 2017). Land cover classification for GHA26 was obtained from the Canada Centre for Remote Sensing (CCRS, 2015) and the availability of each habitat type was assessed. All open canopy land cover types (grasslands, agriculture, etc.) made up a total of 7.4% of the study area. Based on the low availability of open habitat to wolves, we instead chose to examine the influence of the three most available closed canopy habitat types: coniferous, deciduous, and mixed forest. We then calculated the proportion of each habitat type in a 100m buffer around each used and available relocation. Lastly, to incorporate temporal influence on selection, we calculated whether each relocation was recorded during twilight, day, or night using sunset-sunrise data (2014-2019) from the National Research Council of Canada. We defined twilight as the start of civil twilight to one hour after sunrise, as well as from one hour prior to sunset to the end of civil twilight. Day included relocations taken between one hour after sunrise and one hour prior to sunset, and night from 00:00 to civil twilight start and from civil twilight end to 23:59. In our core model, we included each of the habitat proportions, the interaction between step length and turn angle, as well as the interaction between step length and time of day to better understand how wolf habitat selection and movement occurs without the influence of linear features.

### Alternate Model Covariates

In total, we categorized 7 types of linear features in GHA 26. Anthropogenic linear features included primary, secondary, and tertiary roads and three hydroelectric ROWs. Distinguishing these ROWs allowed us to account for potential differences in their influence on wolf selection and movement. The ROWs were classified as following: 1) L5/L47 transmission line, a ROW adjacent to a primary road and hereafter referred to as the ‘paired transmission line’; 2) Line 77 transmission line, an existing ROW that was not paired with any other linear feature; 3) PQ95 transmission line, a relatively new ROW constructed in 2015-16 that typically ranged from 1-3 km from the primary road but occasionally crossed it. We categorized natural linear features such as rivers and shorelines as waterways.

We analyzed two different metrics of linear feature selection (Table 1). First, we calculated the distance, in meters, between each wolf relocation and each linear feature using QGIS3 (QGIS Development Team). Distance to a given linear feature was transformed with the natural logarithm distance +1 to account for the decaying effect of linear features as the distance from them increases (Christina M. Prokopenko et al., 2017). Secondly, we determined if each step crossed each linear feature type, creating a binary variable where 1 indicated crossing and 0 indicated no crossing.

We ran a pair of global models for each linear feature metrics, one using distance to and the other using crossing as measurements of wolf selection for linear features. Each model included the core model as well as the linear feature covariates 1) on their own, 2) interacting with step length, and 3) interacting with the proportion of each habitat type at the end of the step. Following Arce Guillen et al., (2023) we included a random effect for each fixed effect in the models. However, to navigate data processing issues we ran two complementary models, each one with the same fixed effects but only including random effects based for either “habitat selection” or “movement” covariates (Appendix S1).

### Calculating effect sizes

Movement rate was calculated by multiplying the shape and modified scale parameters of the gamma distribution used to generate the available step lengths (Avgar et al. 2016; Prokopenko et al. 2017). As a measure of effect size for selection coefficients, we calculated relative selection strength (RSS) for one location, x1, over another, x2, given the difference in the habitat covariate of interest (Avgar et al. 2017).

## Results

Most wolves selected to be far from linear features with high human activity, i.e., avoided the transmission line adjacent to a primary road (paired) and waterways but strongly selected to be near other linear features with less human activity (P1). At the population level, wolves selected to be close to secondary roads, tertiary roads, and the new transmission line (Figure 2A). There was weak avoidance for primary roads and weak selection for existing transmission lines. As well, responses varied in terms of linear feature crossing where some individual wolves avoided crossing them and others readily crossed these features. Wolves avoided crossing all road types, existing transmission lines and the new transmission line at the population level (Figure 2B).

**Figure 2.**
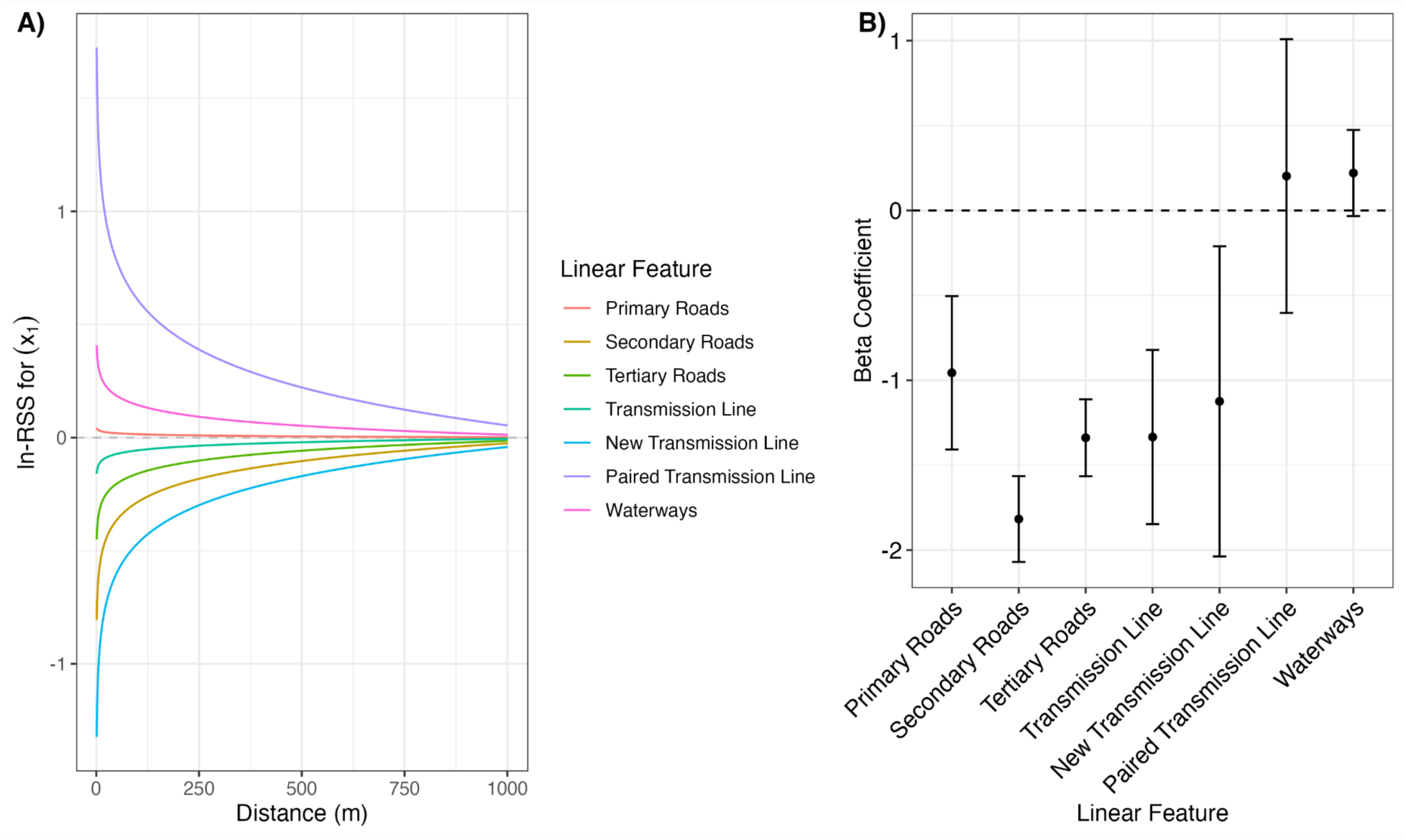
A) Natural log-transformed Relative Selection Strength (ln-RSS) for location x_1_ over another location x_2_ as distance to a linear feature increases at the population level. The dashed line at zero indicates no difference in selection between the two locations. The two locations are identical except for differences in linear feature proximity where x_1_ is 1250 m and x_2_ ranges from 0 to 1000 m. Here, positive values indicate wolves are selecting for x_1_ (i.e., farther distance) and negative values indicate wolves are avoiding x_1_. B) Beta coefficients and 95% confidence interval (CI) indicating the population level (fixed effects) response of wolves to linear feature crossing. Crossing (positive coefficient) or avoidance of crossing (negative coefficient) can be interpreted when the CI does not overlap with zero (dashed line).

Wolves were more likely instead to cross the paired transmission line and had a neutral response to waterways. Wolves increased their step lengths near or crossing linear features, supporting P2 (Figure 3). Wolves exhibited the largest increase in step length when near secondary roads, followed by waterways and primary roads, and only a slight increase in movement for tertiary roads and the pre-existing transmission lines. In contrast, wolves reduced their movement rate when near both the new and paired transmission line types.

**Figure 3.**
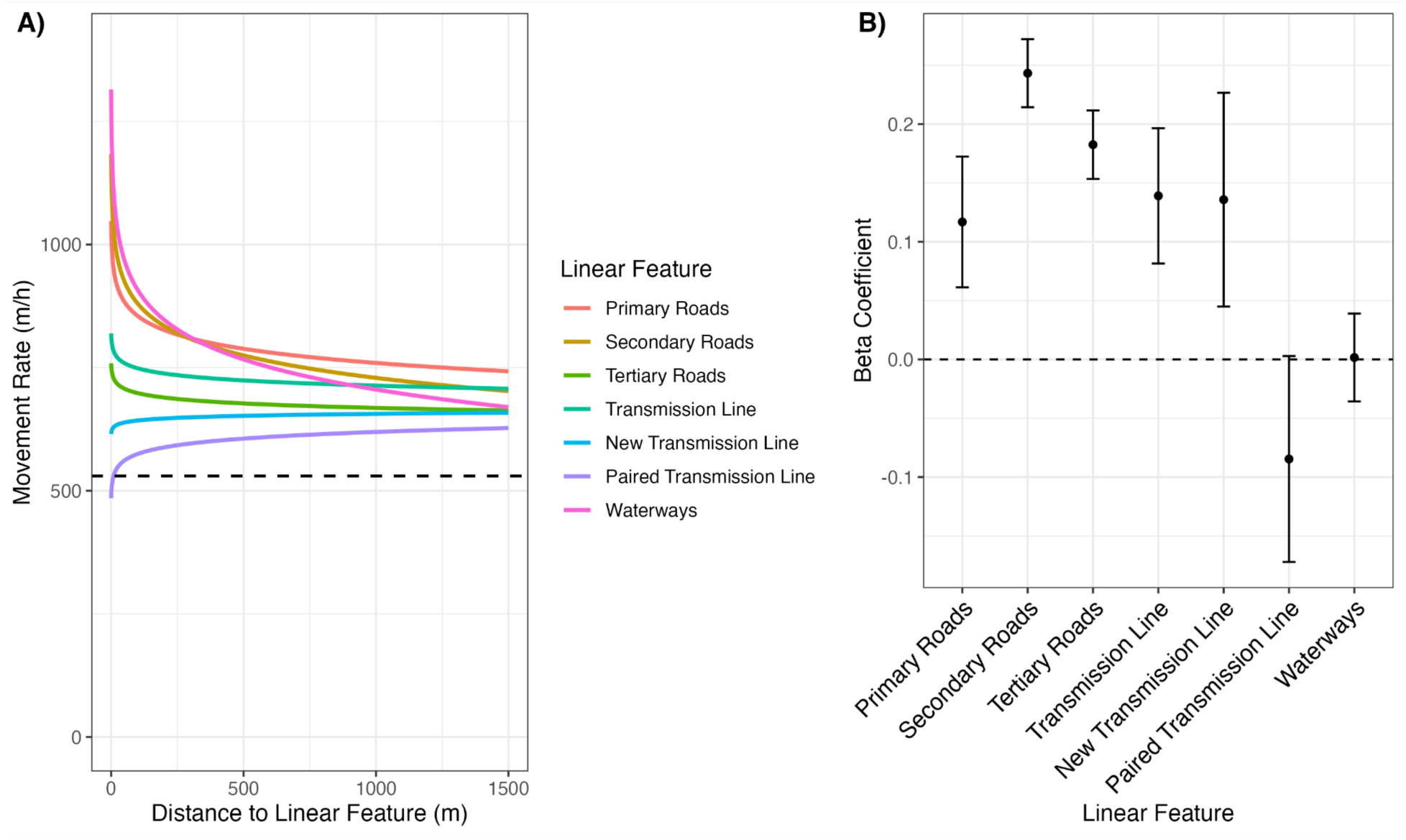
A) Movement response (speed in meters per hour) of wolves as a function of distance to linear features. The dashed line indicates the average step length across the entire population. Speed is calculated from the tentative shape and scale parameters of the gamma distribution of step length and modified by ln-transformed step length coefficients from the model output. B) Shape modifier and 95% confidence interval (CI) indicating the population level movement response of wolves when crossing linear features. Increased movement (positive value) or decreased movement (negative coefficient) when crossing linear features can be interpreted when the CI does not overlap with zero (dashed line).

We found moderate support for our prediction that wolf selection for linear features would increase when in dense forest (P3). In mostly forest habitat, wolves reduced their avoidance of the paired transmission line, particularly when in deciduous or mixed forest habitat (Figure 4). Wolves increased their selection of primary roads when in coniferous forest but increased their avoidance of primary roads when in either deciduous or mixed forest (Figure 4). All forested habitats influenced wolves to switch from avoidance to selection for areas in proximity to waterways. Wolves did not alter their proximity to secondary roads when in any of the forested habitat types but decreased their selection of tertiary roads, most noticeably in deciduous forest (Figure 4). Wolves reduced their selection of the new transmission line in both coniferous and mixed forest, and avoided both this feature and the pre-existing transmission line in high proportions of deciduous forest (Figure 4).

**Figure 4.**
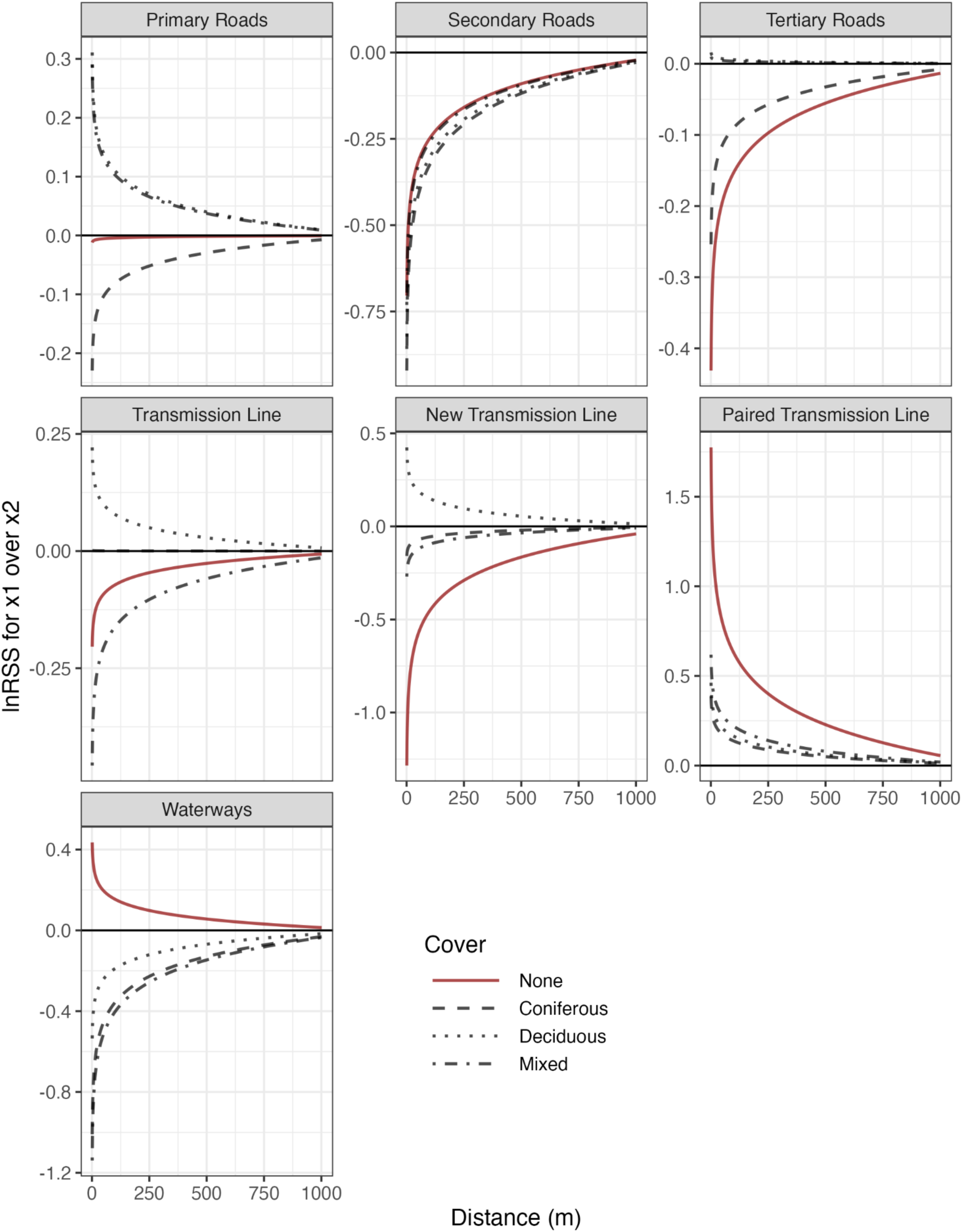
Natural log-transformed Relative Selection Strength (ln-RSS) for location x_1_ over another location x_2_ as distance to a linear feature increases in either no cover (red) or 75 % cover (dashed lines). The solid line at zero indicates no difference in selection between the two locations, which are identical except for differences in linear feature proximity where x_1_ is 1250 m and x_2_ ranges from 0 to 1000 m. Here, positive values indicate wolves are selecting for x_1_ (i.e., farther from the linear feature) and negative values indicate wolves are avoiding x_1_ (i.e., selecting for x_2_).

## Discussion

We studied how a trade-off between risk of human encounter and the reward of movement facilitation influenced wolf use of linear features. Using both proximity and crossing metrics as measures of behaviour, we illustrated that wolves responded to linear features based both on the characteristics of that disturbance and the interaction with the surrounding habitat type. We modelled movement integrated habitat selection within a framework that considered adjacent landscape features and their implicit value or cost. In doing so our method provided a clearer link between the effects of anthropogenically disturbed landscapes and changes in wolf space use behaviour.

Applying a movement and selection framework allowed us to determine if linear features represented risks, barriers, resources, or corridors (Dickie et al., 2020; Prokopenko et al., 2017). When wolves select to be further from linear and increased movement rates when in proximity, these results can indicate wolves perceive areas near these features as high risk (Dickie et al., 2020). Additionally, avoidance of both proximity and crossing of linear features can indicate a barrier effect to wildlife movement (Beyer et al., 2016; Prokopenko et al., 2017). Selection for linear features can indicate increased rewards such as access to resources in proximity to features (e.g., Finnegan et al., 2018), and when combined with increased movement, linear features can be interpreted as travel corridors (Dickie et al., 2020). We found the strongest evidence that combining primary roads with a transmission line created a barrier to movement, and both secondary and tertiary roads were used as travel corridors. We also found moderate support that the new transmission line was generally perceived as a reward, while the pre-existing transmission line and waterways had a more neutral impact on wolf behaviour.

Our results support the hypothesis that wolves avoid linear features with high levels of human activity (Dickie et al., 2017; McKenzie et al., 2012; Whittington et al., 2005). In general wolves exhibited a non-significant response to primary road proximity, however in both mixed and deciduous cover, wolves strongly avoided primary roads but switched their response to selection in coniferous forest. Wolves also avoided crossing primary roads and increased their movement speed when in proximity. Proximity to and crossing of roads increases the risk of wildlife mortality, both in terms of vehicle collisions (Lodé, 2000) and through increased hunting access and success for humans (Gratson and Whitman, 2000). As a result, wildlife species may avoid linear features in response to noise, traffic, or both (Jaeger et al., 2005). The combination of wolves avoiding both proximity to and crossing of features associated with human activity provides evidence that human activity is perceived as risky in this study area. When in dense vegetation such as deciduous or mixed forest, the perceived risk may increase as there is reduced visibility wolves are not able to assess safe crossings, or they may be prevented from escaping human traffic easily. Alternatively, when in coniferous forest, vegetation density is reduced and wolves may instead be able to travel a safe distance from the road while still under the cover of forest canopy. Overall, however, the perception of risk reduces the likelihood of wolves using primary roads as travel corridors or to increase hunting efficiency.

Pairing transmission lines and primary roads elicited a different response from wolves than the response to either feature on their own. Wolves exhibited strong avoidance of areas near the paired transmission line and decreased movement rates indicating a barrier effect to movement. Linear features have been found to act as a barrier to movement in several systems including roads for small forest mammals (Rico et al., 2007) and anuran species (Eigenbrod et al., 2009), and road/river or road/transmission line combinations for moose (Bartzke et al., 2015).

Disturbance associated with linear features can additionally exacerbate functional habitat degradation of the surrounding habitat through avoidance behaviour of perceived risky areas (Dyer et al., 2002; Frair et al., 2008). However, most linear features are not completely impermeable for all species and will allow some movement depending on the species, linear feature width, or traffic volume (Dyer et al., 2002; Frair et al., 2008). Wolves exhibited a neutral response to crossing the paired transmission line, indicating that it is likely semi-permeable to wolf movement. As well, wolves reduced their avoidance of the paired feature in all forested habitats, indicating cover may have a mediating impact on the barrier. As apex predators, the response of wolves could be a signal that these types of features could have connectivity implications for other species as well.

Anthropogenic disturbance alters both predator and prey space use and therefore their interactions (Fahrig, 2003). Prey species are more likely to avoid areas associated with human activity to reduce the risk of predation from both humans and wolves (Kauffman et al., 2007). The reward from linear features for wolves is therefore implied when the risk along a linear feature is reduced. Linear features can increase predator-prey encounter rates through increased predator movement (McKenzie et al., 2012) or search efficiency (Vander Vennen et al., 2016). Wolf selection for and decreased movement in proximity to new transmission line could be indicative of searching behaviour as part of the predation sequence (Dickie et al., 2020). However, wolves reduced selection for or avoided the new transmission line in prey-preferred stands including mixed forest (Ehlers et al., 2014; Kittle et al., 2015). Given the strong selection in low forested areas and relative proximity to the paired transmission line, the new transmission line may provide an alternative to the barrier posed by the paired linear features. Any benefit from selecting transmission lines likely rapidly decreases as vegetation regrows as evidenced from the avoidance of both new and pre-existing transmission lines in deciduous forests.

Wolves selected to be near secondary roads and tertiary roads, which exhibit reduced levels of human activity compared to primary roads. This selection pattern is consistent with previous findings that wolves are more likely to select low-use linear features as the reduction in human-use likely reduces the level of risk to them (Gurarie et al., 2011; Whittington et al., 2005). High forest cover caused wolves to switch their response to strongly select for proximity to waterways indicating there was a benefit to movement when in habitats that posed increased friction to movement. Wolves did not alter selection for secondary roads across habitat types but reduced selection for tertiary roads when in coniferous forest and the switched to avoidance for tertiary roads proximity in dense forested habitat types (mixed and deciduous). Wolves also increased their movement rates when near or crossing low-risk linear features, particularly secondary roads and waterways, indicating wolves may be selecting these features as travel corridors (McKenzie et al., 2012). Tertiary roads in this region are no longer actively maintained and have experienced increased regrowth along these linear features. Regeneration of vegetation would reduce any movement benefit associated with these linear features by increasing friction to movement. As well, waterways were the only linear feature that wolves actively crossed, indicating that natural linear features do not pose a barrier to movement compared to anthropogenic ones.

Monitoring the combined effects of linear features and landscape characteristics demonstrated the risk-reward trade-off greatly influences gray wolf behaviour. The interaction between anthropogenic disturbance and the surrounding landscape can play a mediating role on the tension between human avoidance and resource acquisition. Incorporating the interplay between linear features and the adjacent landscape within a risk-reward framework, as we do here, provides a foundation for the mitigation of human activities. Our work demonstrates that where a linear feature is placed in relation to the surrounding habitat will determine the response of wildlife. High forest cover reduced both the barrier effect of the transmission line paired with primary roads and the movement benefits associated with tertiary roads. This provides evidence that movement corridors such as secondary roads could be actively decommissioned to reduce their impact on increased wolf movement and the potential for increased hunting efficiency. Wolves are apex predators in the boreal food web, and the wolf-ungulate system represents one of the most important trophic cascades for maintaining ecosystem functioning (McLaren and Peterson, 1994). Wildlife typically avoid areas with high anthropogenic activities (Gurarie et al., 2011; James and Stuart-Smith, 2000). However, as human populations, and thus human footprints, expand (Leu et al., 2008; Sih et al., 2011), the spatial separation between humans and wildlife becomes less feasible; instead both species will need to acclimate or adapt to coexist.

## Data Availability

Data and code will be made available upon publication on Zenodo and GitHub.

## Author Contributions

KAK and EVW conceptualized the idea and study design. KAK and DLJD collected the field data. KAK ran the analyses, with input from CMP and JWT, and KAK drafted the article. All authors critically revised the article and gave final approval for publication.

## Acknowledgements

We would like to acknowledge that our research takes place within Treaty 3 and 5 Territory and within the Homeland of the Métis Nation. This work would not be possible without the support of Manitoba Wildlife and Fisheries Branch, and Manitoba Hydro. In particular, we thank K. Leavesley, K Rebizant, D Brannen, D Bulloch, and J Matthewson for their continued support. Thank you to I Lavoie, A Scott, and S Graham for field support. We thank all members of the Wildlife Evolutionary Ecology Lab, including J. Balluffi-Fry, I. Richmond, J. Hogg, J. Kennah, A. Robitaille, J. Aubin, J. Hendrix, Q. Webber, S. Boyle, and L. Newediuk for their reviews. Funding for this study was provided by a CRD Grant to EVW and NSERC CGS-M and CGS-D grants to KAK.

## References

Andrén, H., 1992. Corvid Density and Nest Predation in Relation to Forest Fragmentation: A Landscape Perspective. Ecology 73, 794–804. 10.2307/1940158

Arce Guillen, R., Lindgren, F., Muff, S., Glass, T.W., Breed, G.A., Schlägel, U.E., 2023. Accounting for unobserved spatial variation in step selection analyses of animal movement via spatial random effects. Methods Ecol. Evol. 14, 2639–2653. 10.1111/2041-210X.14208

Avgar, T., Potts, J.R., Lewis, M.A., Boyce, M.S., 2016. Integrated step selection analysis: Bridging the gap between resource selection and animal movement. Methods Ecol. Evol. 7, 619–630. 10.1111/2041-210X.12528

Bartzke, G.S., May, R., Solberg, E.J., Rolandsen, C.M., Røskaft, E., 2015. Differential barrier and corridor effects of power lines, roads and rivers on moose (Alces alces) movements. Ecosphere 6, 1–17. 10.1890/ES14-00278.1

Benson, J.F., Mahoney, P.J., Patterson, B.R., 2015. Spatiotemporal variation in selection of roads influences mortality risk for canids in an unprotected landscape. Oikos 124, 1664–1673. 10.1111/oik.01883

Beyer, H.L., Gurarie, E., Börger, L., Panzacchi, M., Basille, M., Herfindal, I., Van Moorter, B., R. Lele, S., Matthiopoulos, J., 2016. ‘You shall not pass!’: quantifying barrier permeability and proximity avoidance by animals. J. Anim. Ecol. 85, 43–53. 10.1111/1365-2656.12275

Brooks, M. E., Kristensen, K., Benthem, K. J., van Magnusson, A., Berg, C. W., Nielsen, A., Skaug, H. J.., Mächler, M., Bolker, B. M., 2017. glmmTMB Balances Speed and Flexibility Among Packages for Zero-inflated Generalized Linear Mixed Modeling. R J. 9, 378. 10.32614/RJ-2017-066

Coulson, T., Macnulty, D.R., Stahler, D.R., vonHoldt, B., Wayne, R.K., Smith, D.W., 2011. Modeling effects of environmental change on wolf population dynamics, trait evolution, and life history. Science 334, 1275–1278. 10.1126/science.1209441

Courbin, N., Fortin, D., Dussault, C., Courtois, R., 2014. Logging-induced changes in habitat network connectivity shape behavioral interactions in the wolf – caribou – moose system. Ecol. Monogr. 84, 265–285. 10.1890/12-2118.1

DeCesare, N.J., 2012. Separating spatial search and efficiency rates as components of predation risk. Proc. R. Soc. B 279, 4626–4633. 10.1098/rspb.2012.1698

Dickie, M., McNay, S.R., Sutherland, G.D., Cody, M., Avgar, T., 2020. Corridors or risk? Movement along, and use of, linear features varies predictably among large mammal predator and prey species. J. Anim. Ecol. 89, 623–634. 10.1111/1365-2656.13130

Dickie, M., Serrouya, R., McNay, R.S., Boutin, S., 2017. Faster and farther: wolf movement on linear features and implications for hunting behaviour. J. Appl. Ecol. 54, 253–263. 10.1111/1365-2664.12732

Dyer, S.J., O’Neill, J.P., Wasel, S.M., Boutin, S., 2002. Quantifying barrier effects of roads and seismic lines on movements of female woodland caribou in northeastern Alberta. Can. J. Zool. 80, 839–845. 10.1139/Z02-060

Ehlers, L.P.W., Johnson, C.J., Seip, D.R., 2014. Movement ecology of wolves across an industrial landscape supporting threatened populations of woodland caribou. Landsc. Ecol. 29, 451–465. 10.1007/s10980-013-9976-8

Eigenbrod, F., Hecnar, S.J., Fahrig, L., 2009. Quantifying the road-effect zone: Threshold effects of a motorway on anuran populations in Ontario, Canada. Ecol. Soc. 14. 10.5751/ES-02691-140124

Fahrig, L., 2003. Effects of habitat fragmentation on biodiversity FFECTS OF HABITAT FRAGMENTATION ON BIODIVERSITY. Annu. Rev. Ecol. Evol. Syst. 34, 487–515.

Finnegan, L., Pigeon, K.E., Cranston, J., Hebblewhite, M., Musiani, M., Neufeld, L., Schmiegelow, F., Duval, J., Stenhouse, G.B., 2018. Natural regeneration on seismic lines influences movement behaviour of wolves and grizzly bears. PLoS ONE 13. 10.1371/journal.pone.0195480

Fortin, D., Beyer, H.L., Boyce, M.S., Smith, D.W., Duchesne, T., Mao, J.S., 2005. Wolves influence elk movements: Behavior shapes a trophic cascade in Yellowstone National Park. Ecology 86, 1320–1330. 10.1890/04-0953

Frair, J.L., Merrill, E.H., Beyer, H.L., Morales, J.M., 2008. Thresholds in landscape connectivity and mortality risks in response to growing road networkds. J. Appl. Ecol. 45, 1504–1513. 10.1111/j.1365-2664.2007.0

Frid, A., Dill, L., 2002. Human-caused disturbance stimuli as a form of predation risk. Conserv. Ecol. 6, 11.

Gratson, M.W., Whitman, C.L., 2000. Road closures and density and succe of elk hunters in Idaho. Wildl. Soc. Bull. 28, 302–310.

Gurarie, E., Suutarinen, J., Kojola, I., Ovaskainen, O., 2011. Summer movements, predation and habitat use of wolves in human modified boreal forests. Oecologia 165, 891–903. 10.1007/s00442-010-1883-y

Hugie, D.M., Dill, L.M., 1994. Fish and game: a theoretic approach to habitat selection by predators and prey. J. Fish Biol. 45, 151–169. 10.1006/jfbi.1994.1220

Jaeger, J.A.G., Bowman, J., Brennan, J., Fahrig, L., Bert, D., Bouchard, J., Charbonneau, N., Frank, K., Gruber, B., Von Toschanowitz, K.T., 2005. Predicting when animal populations are at risk from roads: An interactive model of road avoidance behavior. Ecol. Model. 185, 329–348. 10.1016/j.ecolmodel.2004.12.015

James, A., Stuart-Smith, A., 2000. Distribution of caribou and wolves in relation to linear corridors. J. Wildl. Manag. 64, 154–159. 10.2307/3802985

Kauffman, M.J., Varley, N., Smith, D.W., Stahler, D.R., MacNulty, D.R., Boyce, M.S., 2007. Landscape heterogeneity shapes predation in a newly restored predator-prey system. Ecol. Lett. 10, 690–700. 10.1111/j.1461-0248.2007.01059.x

Kittle, A.M., Anderson, M., Avgar, T., Baker, J.A., Brown, G.S., Hagens, J., Iwachewski, E., Moffatt, S., Mosser, A., Patterson, B.R., Reid, D.E.B., Rodgers, A.R., Shuter, J., Street, G.M., Thompson, I.D., Vander Vennen, L.M., Fryxell, J.M., 2015. Wolves adapt territory size, not pack size to local habitat quality. J. Anim. Ecol. 84, 1177–1186. 10.1111/1365-2656.12366

Knight, R.L., Kawashima, J.Y., 1993. Responses of raven and red-tailed hawk populations to linear right-of-ways. J. Wildl. Manag. 57, 266–271.

Latham, A.D.M., Latham, M.C., Boyce, M.S., Boutin, S., 2011. Movement responses by wolves to industrial linear features and their effect on woodland caribou in northeastern Alberta. Ecol. Appl. 21, 2854–2865. 10.1890/11-0666.1

Lesmerises, F., Dussault, C., St-Laurent, M.H., 2012. Wolf habitat selection is shaped by human activities in a highly managed boreal forest. For. Ecol. Manag. 276, 125–131. 10.1016/j.foreco.2012.03.025

Leu, M., Hanser, S.E., Knick, S.T., 2008. The Human Footprint in the West : A Large-Scale Analysis of Anthropogenic Impacts. Ecol. Appl. 18, 1119–1139.

Lodé, T., 2000. Effect of a motorway on mortality and isolation of wildlife populations. Ambio 29, 163–166. 10.1579/0044-7447-29.3.163

Luque, G.M., Hochberg, M.E., Holyoak, M., Hossaert, M., Gaill, F., Courchamp, F., 2013. Ecological effects of environmental change. Ecol. Lett. 16, 1–3. 10.1111/ele.12050

Manly, B.F.J., 1974. A Model for Certain Types of Selection Experiments. Biometrics 30, 281. 10.2307/2529649

McKenzie, H.W., Merrill, E.H., Spiteri, R.J., Lewis, M. A., 2012. How linear features alter predator movement and the functional response. Interface Focus 2, 205–216. 10.1098/rsfs.2011.0086

McLaren, B.E., Peterson, R.O., 1994. Wolves, Moose, and Tree Rings on Isle Royale. Science 266, 1555–1558.

Moffatt, S., 2012. Time to Event Modelling: Wolf Search Efficiency in Northern Ontario. University of Guelph.

Muff, S., Signer, J., Fieberg, J., 2020. Accounting for individual-specific variation in habitat-selection studies: Efficient estimation of mixed-effects models using Bayesian or frequentist computation. J. Anim. Ecol. 89, 80–92. 10.1111/1365-2656.13087

Neumann, W., Ericsson, G., Dettki, H., Bunnefeld, N., Keuler, N.S., Helmers, D.P., Radeloff, V.C., 2012. Difference in spatiotemporal patterns of wildlife road-crossings and wildlife-vehicle collisions. Biol. Conserv. 145, 70–78. 10.1016/j.biocon.2011.10.011

Polfus, J.L., Hebblewhite, M., Heinemeyer, K., 2011. Identifying indirect habitat loss and avoidance of human infrastructure by northern mountain woodland caribou. Biol. Conserv. 144, 2637–2646. 10.1016/j.biocon.2011.07.023

Prokopenko, Christina M., Boyce, M.S., Avgar, T., 2017. Extent-dependent habitat selection in a migratory large herbivore: road avoidance across scales. Landsc. Ecol. 32, 313–325. 10.1007/s10980-016-0451-1

Prokopenko, Christina M., Boyce, M.S., Avgar, T., 2017. Characterizing wildlife behavioural responses to roads using integrated step selection analysis. J. Appl. Ecol. 54, 470–479. 10.1111/1365-2664.12768

Rico, A., Kindlmann, P., Sedláček, F., 2007. Barrier effects of roads on movements of small mammals. Folia Zool. 56, 1–12.

Ripple, W.J., Beschta, R.L., 2012. Large predators limit herbivore densities in northern forest ecosystems. Eur. J. Wildl. Res. 58, 733–742. 10.1007/s10344-012-0623-5

Ryall, K.L., Fahrig, L., 2006. Response of predators to loss and fragmentation of prey habitat: A review of theory. Ecology 87, 1086–1093. 10.1890/0012-9658(2006)87 [1086:ROPTLA]2.0.CO;2

Sawaya, M.A., Kalinowski, S.T., Clevenger, A.P., 2014. Genetic connectivity for two bear species at wildlife crossing structures in Banff National Park. Proc. Biol. Sci. 281, 20131705. 10.1098/rspb.2013.1705

Schmitz, O.J., Beckerman, A.P., O’brien, K.M., 1997. Behaviorally mediated trophic cascades: Effects of predation risk on food web interactions. Ecology 78, 1388–1399.

Scrafford, M.A., Avgar, T., Heeres, R., Boyce, M.S., 2018. Roads elicit negative movement and habitat-selection responses by wolverines (Gulo gulo luscus). Behav. Ecol. 29, 534–542. 10.1093/beheco/arx182

Semeniuk, C.A.D., Musiani, M., Birkigt, D.A., Hebblewhite, M., Grindal, S., Marceau, D.J., 2014. Identifying non-independent anthropogenic risks using a behavioral individual-based model. Ecol. Complex. 17, 67–78. 10.1016/j.ecocom.2013.09.004

Signer, J., Fieberg, J., Avgar, T., 2019. Animal movement tools (amt): R package for managing tracking data and conducting habitat selection analyses. Ecol. Evol. 9, 880–890. 10.1002/ece3.4823

Sih, A., Englund, G., Wooster, D., 1998. Emergent impacts of multiple predators on prey. Trends Ecol. Evol. 13, 350–355. 10.1016/S0169-5347(98)01437-2

Sih, A., Ferrari, M.C.O., Harris, D.J., 2011. Evolution and behavioural responses to human-induced rapid environmental change. Evol. Appl. 4, 367–87. 10.1111/j.1752-4571.2010.00166.x

Storm, J.J., Choate, J.R., 2012. Structure and movements of a community of small mammals along a powerline right-of-way in subalpine coniferous forest. Southwest. Nat. 57, 385– 392.

Vander Vennen, L.M., Patterson, B.R., Rodgers, A.R., Moffatt, S., Anderson, M.L., Fryxell, J.M., 2016. Diel movement patterns influence daily variation in wolf kill rates on moose. Funct. Ecol. 1–6. 10.1111/1365-2435.12642

Whittington, J., Clair, C.C.S., Mercer, G., 2005. Spatial responses of wolves to roads and trails in mountain valleys. Ecol. Appl. 15, 543–553. 10.1890/03-5317

