## Supplemental Figures 1-4 for "Movement-integrated habitat selection reveals wolves balance ease of travel with human avoidance in a risk-reward trade-off"

Appendix S1

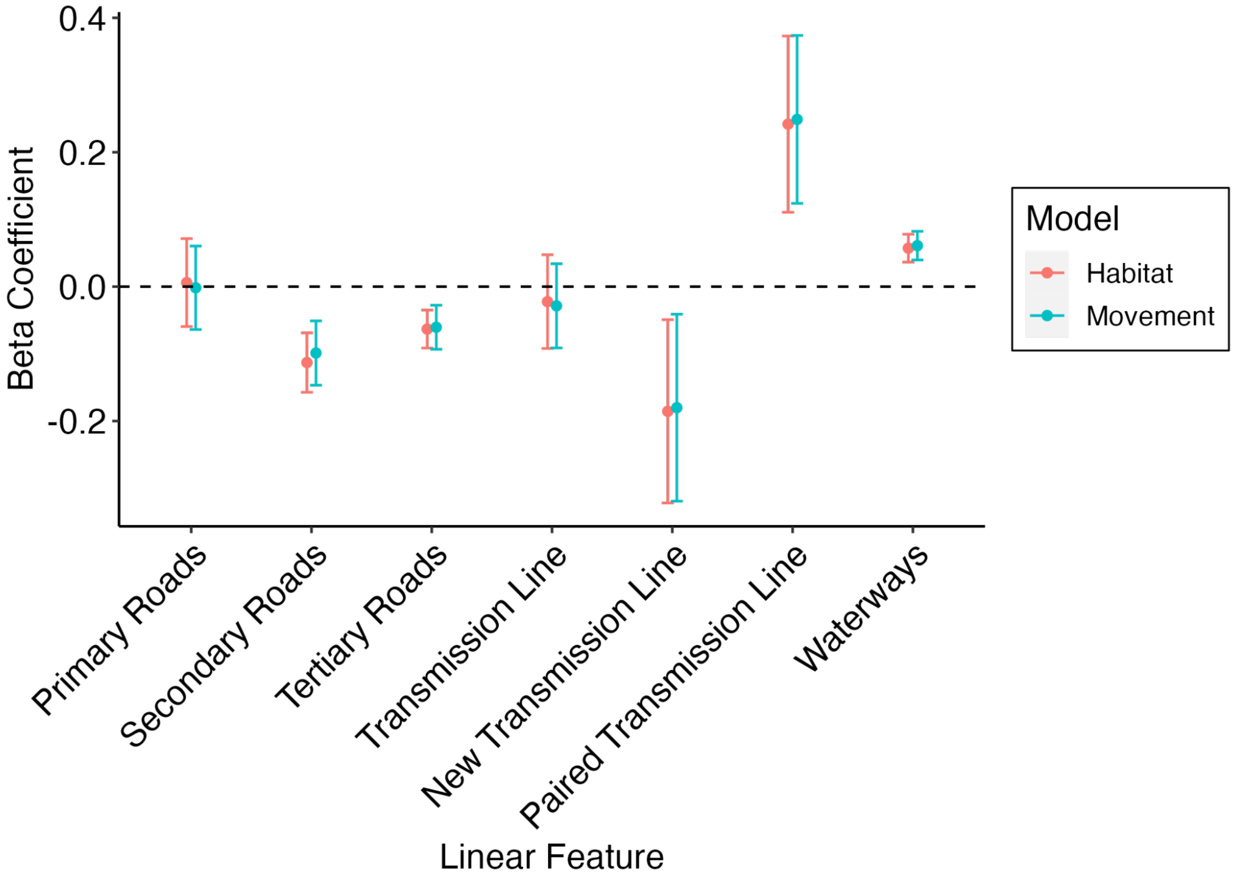


Figure S1. Beta coefficients and 95% confidence interval (CI) indicating the population level response (fixed effects) of wolves (n=45) to linear feature proximity. Comparison of the habitat and movement models indicate that inclusion of model specific random effects did not greatly alter beta estimates. Selection (negative coefficient) or avoidance (positive coefficient) can be interpreted when the CI does not overlap with zero (dashed line).

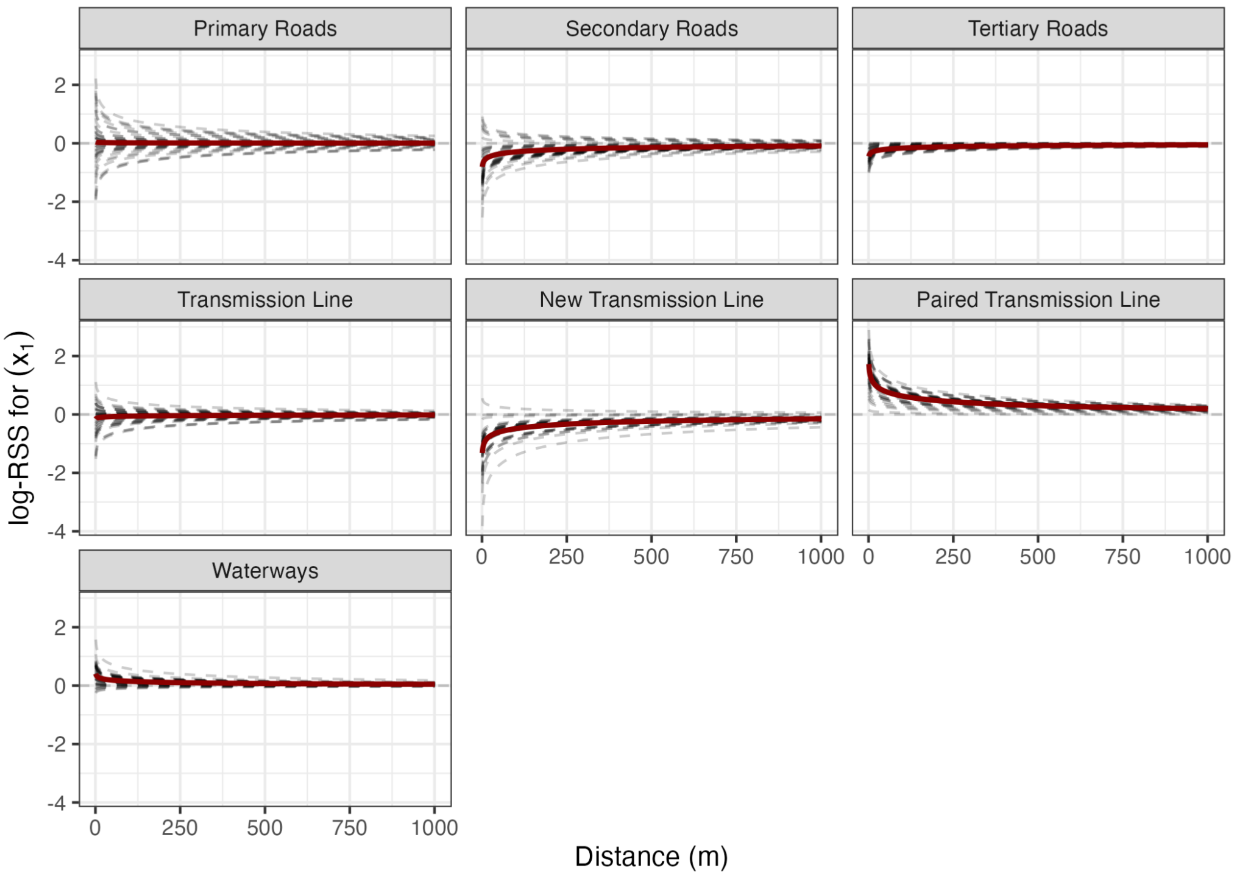


Figure S2. Natural log-transformed Relative Selection Strength (log-RSS) for linear feature proximity for both random (black dashed lines) and fixed (red solid line) effects. The analysis compares selection for location $x_1$ over another location $x_2$ as distance to a linear feature increases. The two locations are identical except for differences in linear feature proximity where $x_1$ is 1250 m and $x_2$ ranges from 0 to 1000 m. Here, positive values indicate wolves are selecting for $x_1$ (i.e., farther distance) and negative values indicate wolves are avoiding $x_1$.

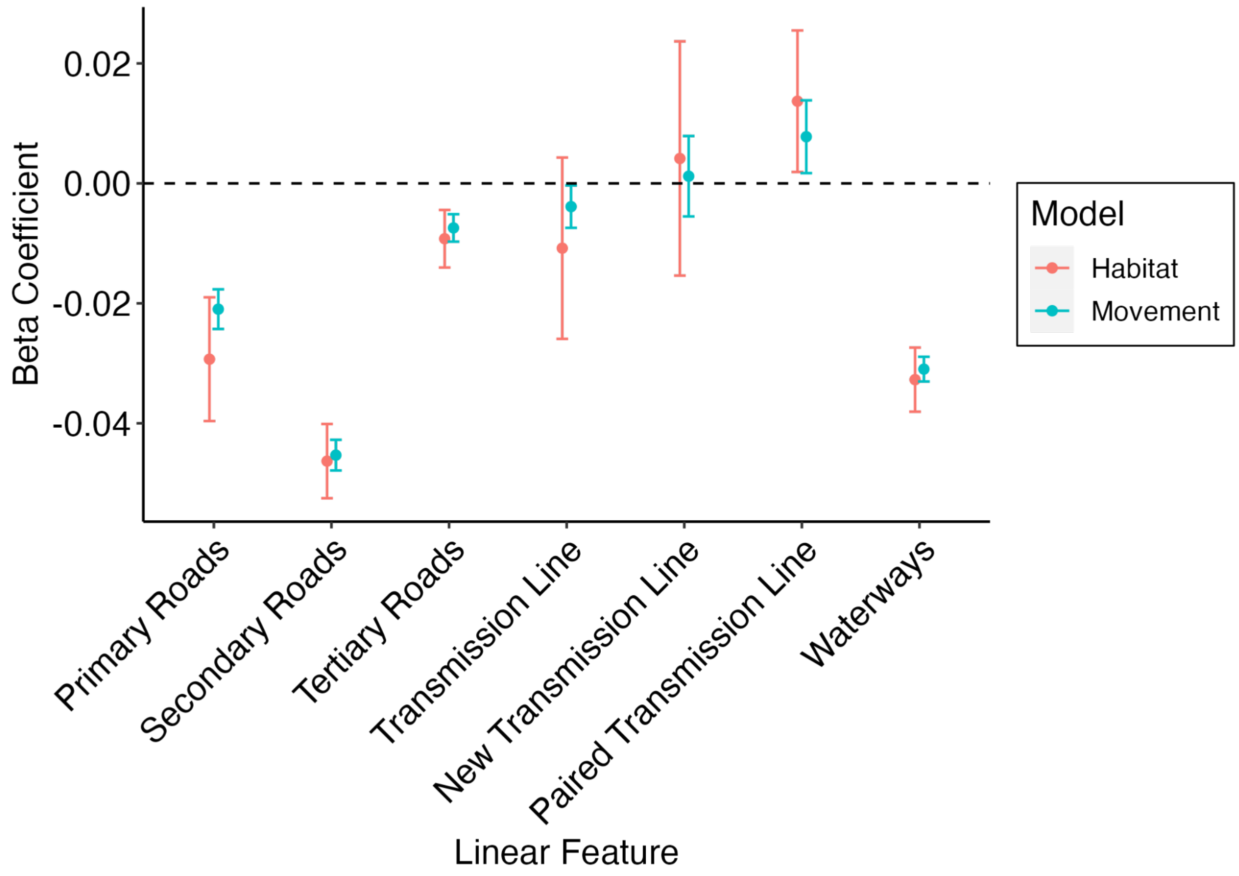


Figure S3. Shape modifier and 95% CI indicating the population level movement response of wolves (n=45) to linear features. Comparison of the habitat and movement models indicate that inclusion of model specific random effects did not greatly alter beta estimates but did improve model fit by decreasing variation around the estimate. Increased step lengths when either near (negative coefficient) or far (positive coefficient) from a linear feature can be interpreted when the CIs do not overlap with zero (dashed line).

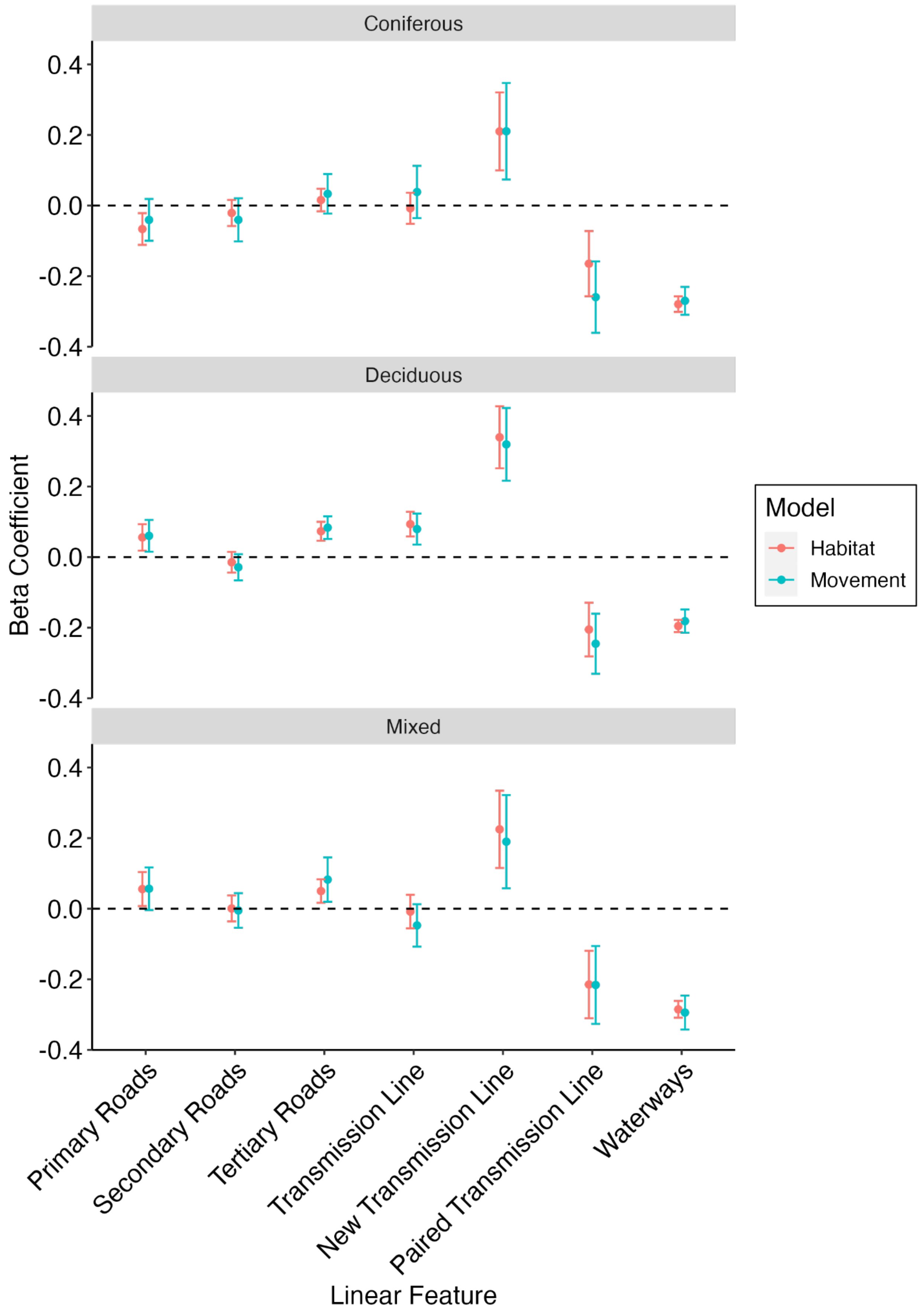

Coniferous
Deciduous
Mixed
Beta Coefficient
0.4
0.2
0.0
-0.2
-0.4
Model
Habitat
Movement
Primary Roads
Secondary Roads
Tertiary Roads
Transmission Line
New Transmission Line
Paired Transmission Line
Waterways
Linear Feature

Figure S4. Beta coefficients and 95% confidence interval (CI) indicating the population level response (fixed effects) of wolves (n=45) to linear feature proximity when interacting with the surrounding habitat type. Comparison of the habitat and movement models indicate that inclusion of model specific random effects did not greatly alter beta estimates. Selection (negative coefficient) or avoidance (positive coefficient) can be interpreted when the CI does not overlap with zero (dashed line).
